# Directing Cholangiocyte Morphogenesis in Natural Biomaterial Scaffolds

**DOI:** 10.1101/2021.04.09.439196

**Authors:** Quinton Smith, Christopher Chen, Sangeeta Bhatia

## Abstract

Patients with Alagille syndrome carry monogenic mutations in the Notch signaling pathway and face complications such as jaundice and cholestasis. Given the presence of intrahepatic ductopenia in these patients, Notch2 receptor signaling has been implicated in driving normal biliary development and downstream branching morphogenesis. As a result, *in vitro* model systems of liver epithelium are needed to further mechanistic insight of biliary tissue assembly. Here, we systematically evaluate primary human intrahepatic cholangiocytes as a candidate population for such a platform and describe conditions that direct their branching morphogenesis. We find that extracellular matrix presentation, coupled with mitogen stimulation, promotes biliary branching in a Notch-dependent manner. These results demonstrate the utility of using 3D scaffolds for mechanistic investigation of cholangiocyte branching and provides a gateway to integrate biliary architecture in additional *in vitro* models of liver tissue.

## INTRODUCTION

The liver is the largest internal organ in the body and is responsible for performing over 500 different vital functions. These tasks include detoxifying drugs, storing nutrients, and producing essential factors such as albumin, clotting proteins, and bile. At a microscopic view, the liver organization consists of repeated hexagonal units termed hepatic lobules. These lobules contain sheets of hepatocytes, flanked by six portal triads, which are comprised of a network of portal veins, hepatic arteries, and intrahepatic bile ducts. In this triad, the vasculature is responsible for the transport of oxygen, nutrients, and clotting factors. The bile ducts, on the other hand, transport hepatocyte-secreted bile acid to the small intestine. Efforts to study liver biology benefit from *in vitro* model systems that recapitulate aspects of the tissue’s native cellular composition and architecture. Existing liver tissue engineering strategies have successfully incorporated human vascular networks with primary hepatocytes *in vitro* ^[1–4]^. However, cell sourcing remains a critical bottleneck in our ability to study human intrahepatic biliary biology. Coupled with this limitation, *in vitro* model systems of the biliary system primarily rely on rodent isolates ^[5]^, immortalized cell lines ^[6,7]^, or adult/pluripotent stem cell-derivatives ^[8–14]^. These populations either do not display mature cholangiocyte marker expression or are limited to forming a non-perfusable structure that lacks the branched architectures found in the native liver.

Immortalized mouse hepatoblasts, with the capacity to differentiate into hepatocytes or cholangiocytes, have been shown to form cystic ductal structures in extracellular matrix (ECM) conditions that contain both laminin rich Matrigel and rat tail collagen I. This finding contrasts with other epithelia, such as Madin-Darby canine kidney cells (MDCK), which only require integrin engagement with collagen I motifs to polarize and expand as cysts ^[15]^. Notably, when mouse hepatoblasts are cultured in collagen I, they form branch-like structures but are unable to polarize. Cystic efficiency in hepatoblast culture is dependent on EGF and HGF stimulation, as well as metalloproteinase and TGFβ activity ^[16]^. Consistent with these findings, immortalized progenitor-like small cholangiocytes derived from mice cannot spread in collagen I matrices or form cystic structures in Matrigel but require decellularized liver ECM to undergo branching morphogenesis ^[17]^. To reduce the complexity of xeno-derived matrices, synthetic hydrogel scaffolds can be engineered with specific material properties such as stiffness, porosity, and adhesion densities, permitting systematic decoupling of physicochemical cues necessary for tissue homeostasis and morphogenesis. For example, immortalized normal rat intrahepatic cholangiocytes (NRCs) ^[7]^ encapsulated in polyethylene glycol (PEG) pre-polymers that are functionalized with fibronectin-derived RGD binding motifs can expand as cysts in a stiffness-dependent manner. Soft (0.5 kilopascal) hydrogel matrices lead to frequent cyst formation, and increased RGD concentrations encourage multi-lumenal features, but interconnected branched epithelial structures could not be generated ^[18]^. Furthermore, a variety of approaches have been used to build perfusable biliary tubes, but the resulting channels lack hierarchical structure, and are composed of rodent-derived cholangiocytes ^[19,20]^. To date, the vast majority of studies in this field have relied on rodent-derived cellular material, and there are many examples in the liver tissue engineering field in which findings obtained using mouse and rat cells do not correlate with the outcomes obtained with human samples ^[21–25]^. While the advent of immortalized human biliary cell lines can help to reduce these variances, they present their own limitations, including a heavy mutational burden that can lead to clonal variability from the original source and transformation from the natural phenotype as in the case with hepatocytes ^[26]^. Motivated both by these advances and the remaining progress gaps, we sought to build upon existing biliary platforms and investigated the potential to fabricate branched human cholangiocyte networks, alongside cholangiocyte-lined channels.

To this end, we characterized commercially available adult-derived primary human cholangiocytes and investigated their branching potential in 3D culture conditions. First, we performed cellular profiling with biliary-specific surface markers and measured tissue-specific enzymatic activity. After validating cholangiocyte-like identity of these isolates, we conducted 3D culture experiments in natural biomaterial scaffolds containing varying concentrations of mitogens relevant to liver development and regeneration. From these studies, we found that growth factor cues and the extracellular matrix coax *in vitro* biliary network assembly in a Notch signaling-dependent manner^[1,27,28]^. In addition, we demonstrate that branching architectures of this human cholangiocyte-like population are also supported in an engineered microfluidic platform that has been used previously as an organ-on-chip model system^[1,27,28]^. This finding highlights the potential to harness these primary human cells in this perfusable platform for future investigations of cholangiocyte functionality including permeability, shear stress response, and transport of bile fluid components such as cholic/chenodeoxycholic acids or xenobiotics. Collectively, our combined approaches reinforce the role of EGF stimulation and Notch signaling in biliary morphogenesis.

## RESULTS

### Primary Intrahepatic Cholangiocytes Maintain Functional Marker Expression *In Vitro*

Sourcing human primary cholangiocytes remains a challenge due to their intrahepatic localization, however pluripotent stem cell derivatives ^[10,29–31]^ or Lgr5+ enriched adult duct progenitor populations acquired from biopsied samples ^[9]^, have been routinely used as model systems to date. While these sources permit the expansion of cholangiocyte-like cells as cystic organoids, their maintenance requires administration of a complex chemical milieu that maintains a stem-like state and does not promote the self-assembly of physiologically relevant branched architectures. Here, we appraised morphological and functional features of commercially available, adult intrahepatic biliary epithelial cells (IHCs) using a combination of tools including gene expression analysis, flow cytometry and immunofluorescence staining (Fig. 1A). Phenotypic assessment was performed on thawed biliary cells that were expanded on collagen type I coated plates for up to 4 passages. We first performed flow cytometry analysis to assess the homogeneity of these cell populations, as well as the degree of mature protein marker expression. As expected, the majority of IHCs expressed biliary markers SRY-related HMG transcription factor 9 (SOX9), cytokeratin 7 (CK-7), cystic fibrosis transmembrane conductance regulator (CFTR), and a subset were positive for epithelial cell adhesion molecule expression (EpCAM). In contrast with stem cell-derived sources, IHCs expressed low levels of the mature marker somatostatin receptor 2 (SSRT-2), a mediator of hormonal signals during digestion, and did not exhibit alpha-fetoprotein (AFP), a common marker expressed by progenitors (Fig. 1B).

**Figure 1.**
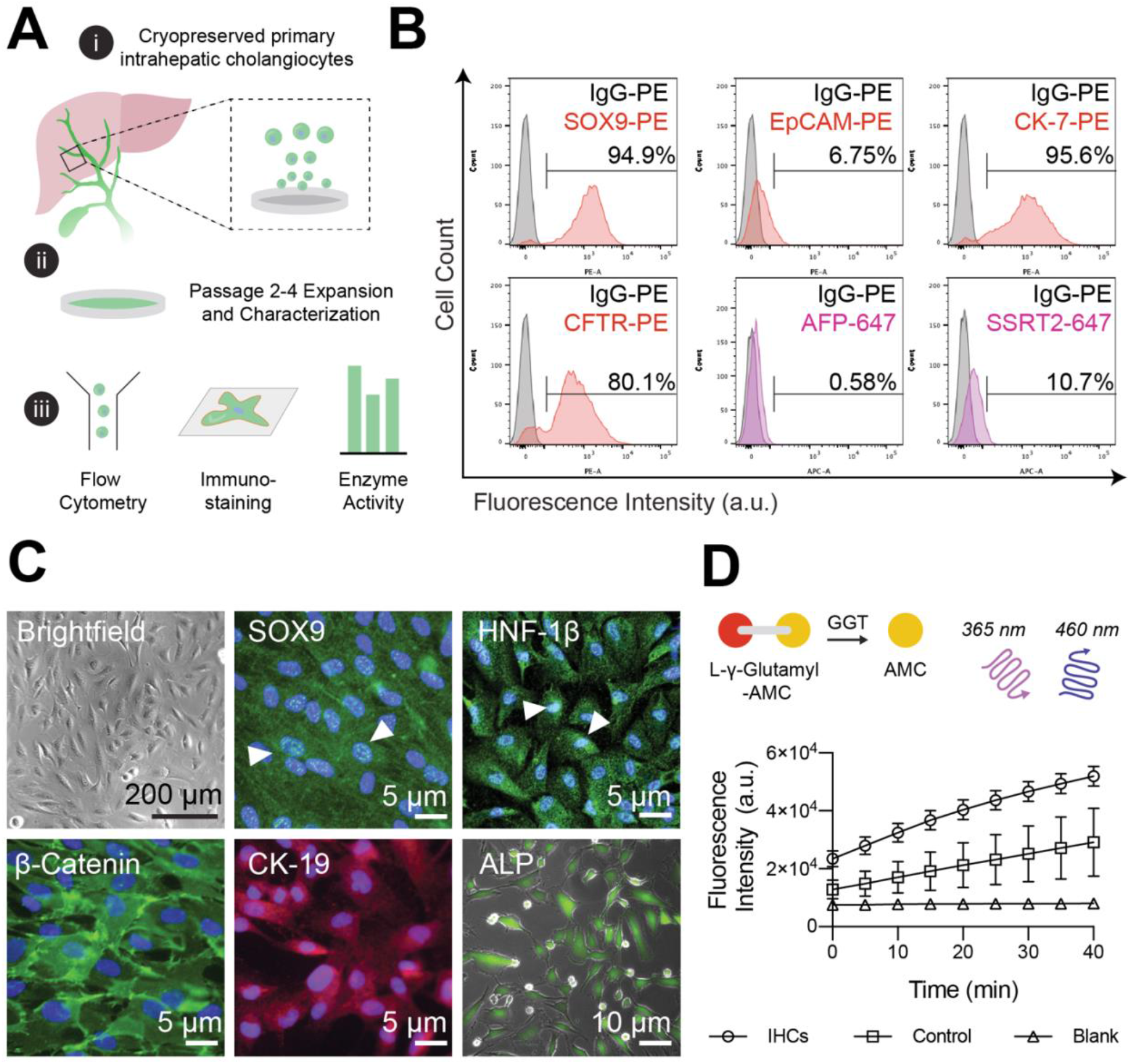
Characterization of primary intrahepatic cholangiocytes. **(A)** (i) Schematic describing the approach to characterize adult, cryopreserved primary human intrahepatic cholangiocyte (IHCs); (ii) IHCs were expanded on collagen-coated plates and used for up to four passages; (iii) Passage 2 IHCs were characterized with a combination of flow cytometry analysis, immunofluorescence staining, and enzymatic activity assays. **(B)** Representative flow cytometry histogram plots showing SOX9, EpCAM, CK-7, CFTR, AFP and SSRT2 expression compared to IgG isotype controls (in gray; [n = 3]). **(C)** Representative brightfield and epi-fluorescence images of IHCs cultured on collagen-coated substrates (n = 3 biological replicates). IHCs show nuclear localization of SOX9 and HNF-1β expression (green signal overlapping with nuclear DAPI stain in blue; white arrows), and positive cytoplasmic expression of β-catenin (green) and CK-19 (red). IHCs were incubated with a live alkaline phosphatase (ALP) stain for 30 minutes, washed with serum-free media and imaged with fluorescence microscopy (Nuclei were stained with DAPI). **(D)** γ-glutamyl transferase activity (GGT) measured using a colorimetric assay where time course of fluorescence intensities is shown, denoting the liberation of 7-Amino-4-Methyl Coumarin (AMC) from a γ-glutamyl quenched substrate. Graphs represent mean and standard deviation from technical triplicates with blank and GGT positive controls.

We observed that cultured IHCs exhibited polygonal morphology, with visible cell-cell adhesions (Fig. 1C). While IHCs initially grew as distinct patches with regular borders, after multiple passages they began to exhibit elongated and spindle-like morphologies, indicative of epithelial to mesenchymal transition, and acquired some fibroblastic features such as fibronectin deposition while maintaining a degree of junctional marker expression (Supplementary Fig. S1). Based on this observation, IHCs were not used for functional studies beyond five passages. Immunofluorescence microscopy was used to visualize the localization and expression of proteins common to epithelial and cholangiocyte identity, namely membrane bound cell-cell adhesion marker β-Catenin, cytoplasmic cytokeratin 19 (CK-19) and nuclear biliary markers hepatic nuclear factor - 1β (HNF-1β) and SOX9 (Fig. 1C). Zinc metalloenzymes alkaline phosphatase (ALP) and γ-glutamyl transpeptidase (GGT) are present in nearly all tissues but are enriched in biliary epithelium *in vivo* and we observed that this population of IHCs demonstrate these functional features (Fig. 1C, D; Supplementary Fig. S1). Through the collective analysis of protein expression and enzymatic activity, we presume that the IHC population consists mainly of large rather than small cholangiocytes, evidenced by CFTR expression and GGT and ALP activity ^[32]^.

### Composite Extracellular Matrices Promote Cholangiocyte Branching in 3D Culture

After validating that IHCs exhibit a collective set of phenotypic and functional traits, we proceeded to leverage insight from *in vivo* developmental studies to assay whether these cholangiocyte-like cells can self-assemble into biliary networks *in vitro*. We hypothesized that IHCs would have the ability to form interconnected 3D branched network structures within a native extracellular niche, given the appropriate introduction of matrix and chemical cues. To this end, we first fluorescently labeled IHCs using a puromycin-selective lentiviral red fluorescent protein (RFP) system, designed to visualize live F-actin expression. Next, after selection via antibiotic resistance, we expanded and encapsulated the resulting Life-Act RFP-IHCs at a density of 1×10^6^ cells/mL in either Matrigel or Matrigel/ collagen type 1 blends. Four days post encapsulation, we fixed and labeled the nuclei of resulting structures, imaged using confocal microscopy, and performed image analysis on z-plane maximum intensity projections. To evaluate the resulting morphological features, we used a computer algorithm to segment image attributes and quantified network coverage, branch points, and structure lengths (Fig. 2A).

**Figure 2.**
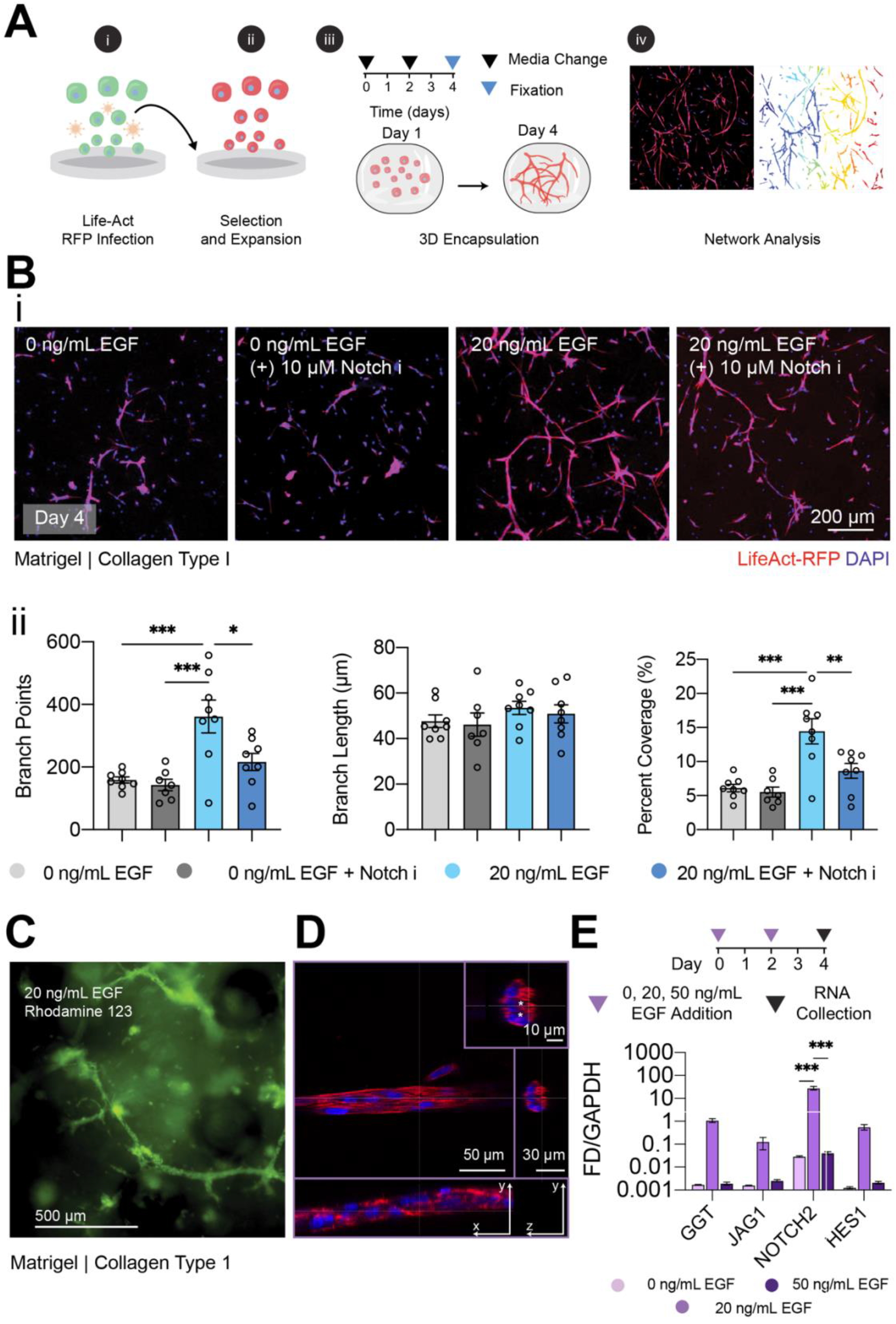
Composite Matrigel and Collagen Type I Gels Support Cholangiocyte Branching Morphogenesis. **(A)** Experimental schematic to probe the capacity for primary human intrahepatic cholangiocytes to undergo branching morphogenesis: (i) Fluorescence tagging of IHCs by rLVUbi-LifeAct-TagRFP cytoskeletal labeling; (ii) A homogenous population of IHCs stable for red fluorescence protein (RFP) expression of F-actin was generated after puromycin selection and expanded in cholangiocyte media; (iii) Resulting populations were encapsulated in Matrigel/collagen type 1 gels for four days, and (iv) features of branching morphologies were measured from maximum intensity z-projections of confocal images using a segmentation algorithm. **(B)** (i) Representative images of LifeAct RFP-IHCs encapsulated in Matrigel/Collagen I blends, cultured with and without EGF or Notch inhibition (10 µM L,685,458). (ii) Individual data points of quantified branch length, points, and network percent coverage. P-values were obtained using One-Way ANOVA Tukey’s hypothesis testing. Representative images generated from at least 7 independent fields of view from 3 biological replicate experiments. At least 20 segmented features were analyzed per field of view. **(C)** Functional uptake of Rhodamine 123 after 4 days of IHC culture in Matrigel/collagen type 1 blends containing 20 ng/mL of EGF. **(D)** F-actin stain of cholangiocytes grown in Matrigel/collagen type 1 composites after 4 days with 20 ng/mL EGF stimulation. White asterisks in the cross-section indicate lumen within the branched structures **(E)** mRNA expression levels of *GGT* and Notch signaling genes measured via RT-qPCR for IHCs four days post 3D culture. Bar graphs show internal triplicate measurements from the pooled collection of 5 biological replicate gels. P-values were obtained using Two-Way ANOVA Tukey’s hypothesis testing. P < 0.033 (*), P< 0.002 (**), P <0.001 (***). All data represented as mean ± SEM.

IHCs that were encapsulated in growth factor-reduced Matrigel were incubated with or without EGF stimulation and exposed to conditions with L-685,458, a potent and selective γ-secretase inhibitor that blocks Notch transcriptional activity ^[33]^. IHCs cultured in pure Matrigel scaffolds without EGF stimulation produced limited sprouting and formed large aggregate structures, with multiple branched features extending from the clustered core. With Notch inhibition, cell aggregation diminished, indicative of increased matrix interaction compared to homotypic cell-cell interactions. However, multi-cellular branched features were not identified (Supplementary Fig. S2). Sprouting behavior in Matrigel improved in the presence of 20 ng/mL EGF, demonstrating the role of mitogen stimulation on biliary branching in laminin rich matrices (Supplementary Fig. S2).

We hypothesized that the addition of fibrillar architecture within Matrigel could promote biliary sprouting behavior. Therefore, we created composite gels consisting of Matrigel and 3.0 mg/mL rat tail collagen type I and assessed IHC sprouting behavior under EGF stimulation and Notch inhibition. While there were no significant differences in the number of branch points, branch length, or percent coverage of IHCs cultured in 3D composite gels with 0 ng/mL EGF with or without Notch inhibition, we found that the addition of 20 ng/mL EGF had a significant impact on the potential to form interconnected branched features. Notably, we observed an increase in the density and number of observed branched points (Fig. 2B i, ii). We also evaluated the combinatorial role of co-administering hepatocyte growth factor (HGF) with EGF on IHC branching potential, as they both have been implicated in inducing ductal morphogenesis during development (Supplemental Fig. S3A). We found that HGF alone can also support IHC sprouting, but the combination with HGF and EGF leads to densely interconnected structures. Again, when Notch inhibition is introduced, the additive effects of HGF and EGF are abrogated, implicating a strong role for Notch signaling in branching morphogenesis (Supplemental Fig. S3B, C).

We next appraised network functionality through analysis of ATP-dependent flux. The multidrug resistance 1 (MDR1) P-glycoprotein protects cholangiocytes from toxic cationic agents present in hepatocyte-secreted bile acid, including xenobiotic substances or drugs. Rhodamine 123 is a fluorescent tracer dye and substrate of MDR1 and is actively transported into the lumen of biliary epithelial cells. 3D incubation of Rhodamine 123 in 20 ng/mL EGF-stimulated networks led to secretory functionality, with a lumenal influx of the fluorescent substrate (Fig. 2C). We were also able to identify lumen within the branching cholangiocytes using high-resolution confocal microscopy (Fig. 2D). For a controlled evaluation of the role of growth factor presentation during IHC branching, we chose to specifically look at EGF stimulation in 3D culture conditions, but also compared these effects to IHCs cultured as 2D monolayers (Supplementary Figure S4). To elucidate the effects of EGF on known Notch signaling targets, IHC transcript levels were measured in 3D composite gels cultured with 0, 20, or 50 ng/mL EGF. Our analysis revealed that EGF stimulation affected *GGT, JAG1, NOTCH2*, and *HES1* mRNA expression levels in a dose-dependent manner and elicited a significant increase in *NOTCH2* gene expression with 20 ng/mL EGF stimulation (Fig. 2E).

### EGF Stimulation Enhances Notch Signaling During Cholangiocyte Branching

We engineered Notch2-deficient cells using CRISPR/Cas9 (Clustered Regularly Interspaced Short-Palindromic Repeats/ CRISPR associated protein 9) mediated deletion to further validate EGF’s role in Notch signal transduction and IHC branching potential. In brief, two guide RNAs (gRNAs; Scramble control, and Notch2) were cloned into puromycin-sensitive lentiviral CRISPR/Cas9 vectors and packaged with HEK293 cells. The resulting complexed particles were used to infect freshly thawed IHCs (Fig. 3A). Relative protein levels of the Notch2 intracellular domain (Notch2-ICD) were confirmed via western blot analysis of cell lysates, showing decreased expression in Notch2 knockdown (KD) compared to scrambled control cells (Fig. 3B). These changes were consistent at the mRNA level (Fig. 3C), reflecting a broad uptake of the CRISPR-mediated deletion, despite some residual, Notch2-intact cells remaining in the population. Finally, cells from the control and Notch2 depleted populations were encapsulated in Matrigel/collagen type 1 composite gels with 20 ng/mL EGF stimulation. After 4 days, the Notch2-depleted population exhibited dramatically blunted sprouting potential, relative to the scrambled control cells (Fig. 3D). These results confirm the role of EGF in the Notch signaling axis in IHCs, which presumably mediates branching morphogenesis in composite natural scaffolds.

**Figure 3.**
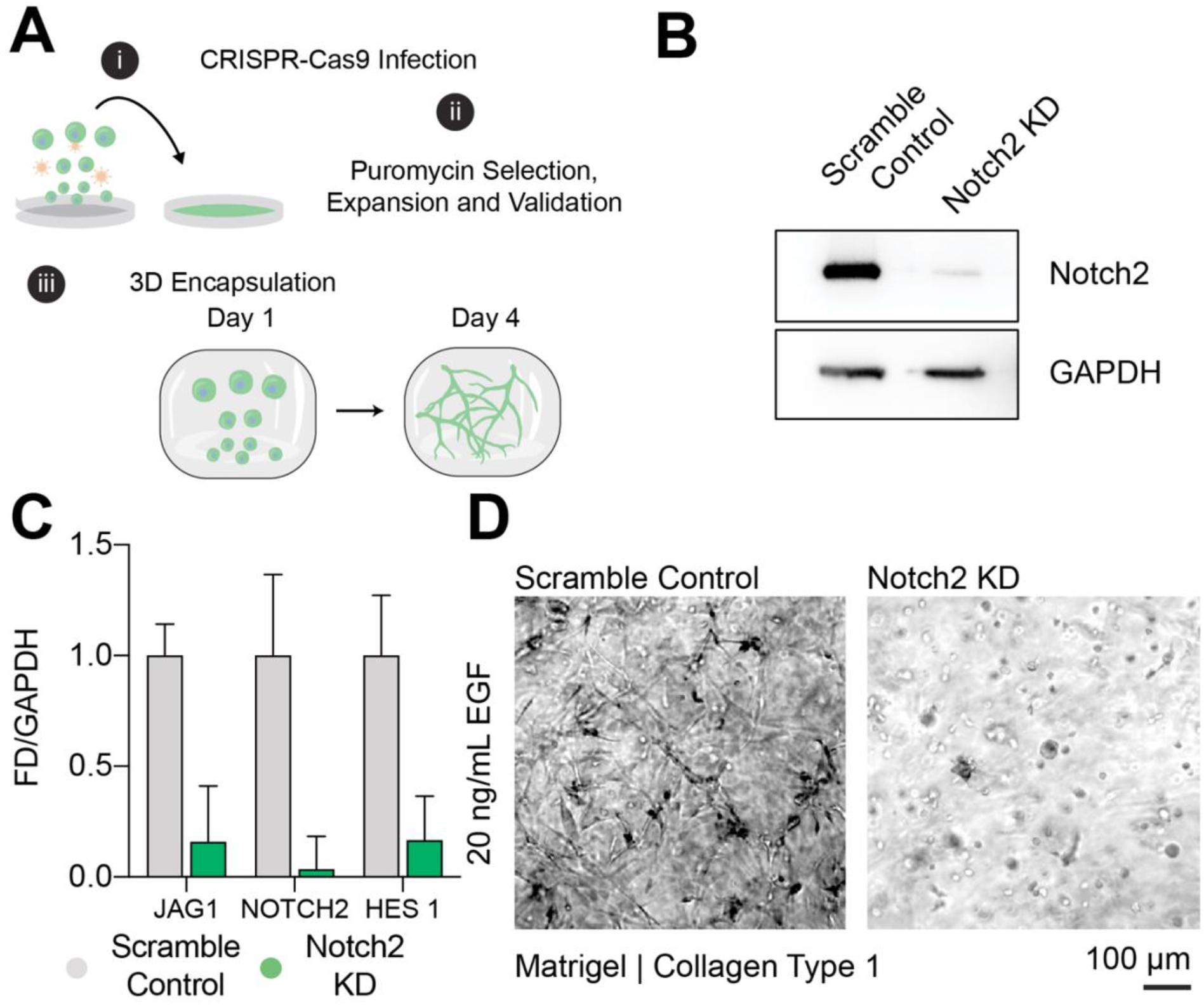
CRISPR-Cas9 Mediated Notch2 Knockdown Ablates Cholangiocyte Branching Morphogenesis in Hydrogel Blends. **(A)** Schematic of the workflow to test the impact of Notch2 knockdown in cholangiocyte branching morphogenesis. **(B)** Notch 2 intracellular domain (Notch 2 – ICD) downregulated protein expression in Notch 2 knockdown (KD) IHCs compared to scramble control cells. **(C)** Comparison of mRNA expression between scramble control and Notch2 KD cells. Notch2 KD cells show downregulation of *JAG1, NOTCH2*, and *HES1* gene expression compared to scramble controls (n = 2). **(D)** Brightfield images of scramble control and Notch2 KD cells grown in Matrigel/ collagen I hydrogel blends after four days. Notch2 KD cholangiocytes show blunted branching compared to control cells (n=3, triplicate gels).

### Intrahepatic Biliary Tree on a Microfluidic Chip

To demonstrate IHC sprouting potential in a system amenable to flow, which allows the study of biliary phenotypes in a dynamic, physiologically relevant microenvironment, we leveraged a previously developed microfluidic platform ^[27,34]^. In brief, the polydimethylsiloxane (PDMS)-based device contains guide features when bonded on a coverslip, allowing for the insertion of parallel needles (300 µm in diameter, spaced 1 mm apart). Subsequent to needle insertion, prepolymer solutions with or without cells can be introduced within the device. After the polymer has crosslinked, needle removal leaves open structures that can be seeded with cells, allowing for bulk cellular self-assembly and prefabricated vessel-shaped structures (Fig. 4A). We first leveraged insight from our bulk 3D culture experiments, and mixed IHCs in pre-polymerized composite gels into the device. Following polymerization and needle removal, we seeded additional IHCs into the lumen of the channels via a pressure differential and cultured the devices under gravity-driven perfusion using a rocker platform. After monitoring RFP expression over the course of 6 days, we found that the vessel structures collapsed into cords, blunting channel access for perfusion, but permitted the assembly of branching architectures in the bulk of the device (Fig. 4B). When comparing this observed phenotype to microfluidic culture of normal rat cholangiocytes (NRCs), we found that Matrigel/collagen blends supported both bulk cystic morphogenesis and sustained polarized cell-laden channels as previously described (Supplementary Figure S5)^[19]^. NRC culture inside the fluidic channels phenocopies the epithelial polarization we expect in normal biliary epithelium, including strong junctional marker expression. However, to maintain patent vessel architecture with the IHCs, we explored the use of alternative biomaterial scaffolds. We found that fibrin, a sticky clotting agent comprised of fibrinogen and thrombin, enhanced biomaterial adhesion to the glass and PDMS features of the device, sustaining both patent cholangiocyte-lined ducts as well as dense, interconnected branched features (Fig. 4C). Collectively, these results demonstrate the capacity to form a biliary tree on a microfluidic chip that is amenable to the introduction of flow.

**Figure 4.**
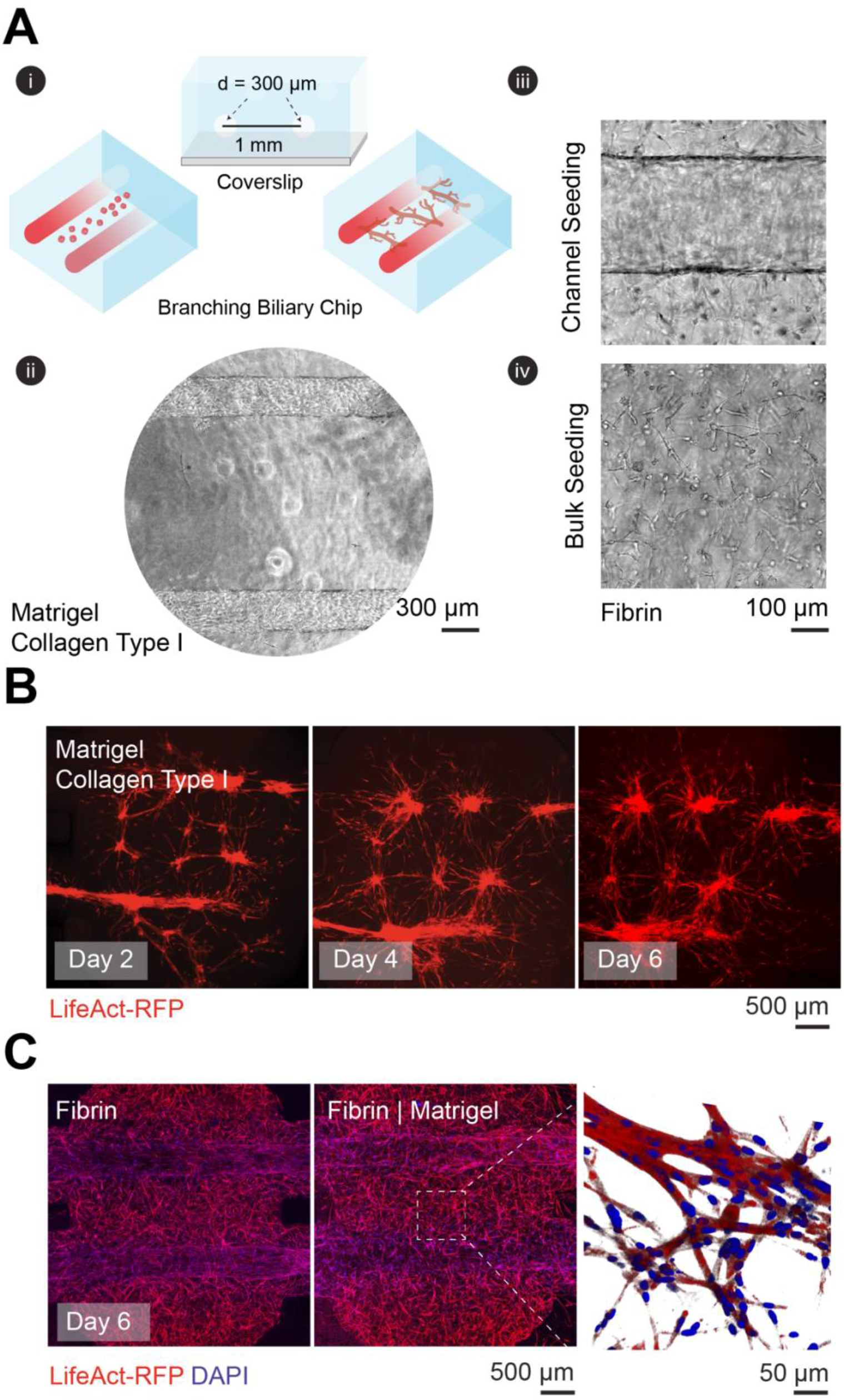
Hierarchical Intrahepatic Biliary Duct on a Chip. **(A)** (i) Device schematic and representative brightfield images, showing top and cross-sectional views of a dual channel microfluidic platform; (ii) encased within the biomaterial scaffolds are two parallel 300 µm open lumenal structures spaced 1 mm apart; (iii) the arrangement provides capacity to seed IHCs and flow media in the patent channels; (iv) and encapsulate IHCs in the bulk compartment of the device. **(B)** Representative time course images of LifeAct RFP-IHCs grown in microfluidic device with 20 ng/mL EGF, showing collapse of open structures after six days, but anastomosis between bulk networks with channel structures. **(C)** Representative maximum intensity projection images of LifeAct RFP-IHCs cultured for 6 days with 20 ng/mL EGF in fibrin scaffolds, showing maintenance of open cell-laden channels compared to Matrigel/collagen type 1 scaffolds (Nuclei stained with DAPI).

## DISCUSSION

Here we describe a pipeline for characterizing primary human biliary isolates and appraised their functional and morphogenic potential under controlled *in vitro* microenvironments. We specifically show that extracellular matrix composition, growth factor presentation, and Notch activity elicit the self-assembly of branched epithelial structures that mimic native biliary tissue. The resulting structures maintain cholangiocyte-specific function and are able to transport multiple drug resistance protein substrates. Furthermore, we successfully combined top-down and bottom-up approaches to engineer large (300 µm) cholangiocyte-lined channels as well as smaller self-assembled branched cholangiocyte networks in a microfluidic platform. We envision that this engineered ‘chip’ format can be leveraged for future evaluation of shear stress, permeability, and spatially organized co-cultures. Collectively, these results reinforce approaches that can be used to study cell-cell and cell-matrix during tissue morphogenesis and provides a strategic framework for future integration of biliary epithelium in existing engineered liver platforms ^[35,36]^.

Notch signaling is an evolutionarily conserved intracellular pathway that regulates many aspects of embryonic development including cell-fate specification and morphogenesis. Consequently, dysregulation of Notch signaling can lead to an array of developmental disorders that affect many tissues including the liver. Mammalian cells contain four different single pass transmembrane Notch receptors (NOTCH 1-4). Upon ligand engagement by neighboring Jagged or Delta-like protein expressing cells, the extracellular domain of the Notch receptor undergoes proteolytic cleavage by an ADAM metalloprotease. Following this reaction, cleavage of the Notch intracellular domain (NICD) by γ-secretase, liberates the protein to translocate to the nucleus where it interacts with the recombination signal binding protein for immunoglobulin kappa J (RBPJ) transcription factor. This association promotes downstream expression of hairy and enhancer of split-1 (HES1) which acts to confer instructions to neighboring cells during embryonic patterning. Notch signaling is essential to bile duct specification and subsequent tubule formation ^[37]^, and patients with Alagille syndrome, contain inherited mutations in either the Notch2 receptor or Jag1 ligand resulting in cholestasis from ductal paucity^[38]^. Kitade et al. show mouse derived bipotential hepatic progenitor cells (HPCs) undergo cholangiocyte specification and branching morphogenesis in Matrigel/collagen I gels. This branching is mediated by EGFR and MET stimulation using EGF and HGF media supplementation, respectively. *EGFR* null HPCs are unable to undergo branching morphogenesis or acquire biliary marker expression upon growth factor addition ^[39]^. Furthermore, *EGFR* competent HPCs with Notch deletion fail to differentiate or branch with MET/EGFR activation. In line with these results, our human *in vitro* system corroborates the crosstalk between Notch/EGF (EGFR) signal transduction in biliary morphogenesis ^[37,40]^.

We show that inhibition of γ-secretase by L-685,458 or knockdown of Notch2 via CRISPR/Cas9, leads to reduced branching morphogenesis *in vitro* and abrogates transcriptional activation of *HES1*. Furthermore, we find that EGF stimulation during branching morphogenesis is correlated with increased Notch signal transduction, evidenced by the upregulation of *HES1*. We find that this transcriptional regulation of Notch activity is specific to 3D culture conditions. Whether this phenotype is regulated by ECM stiffness has yet to be elucidated, but mechanical forces have been demonstrated to regulate YAP/TAZ control of Notch activation in epidermal stem cells ^[41]^. Additionally, bi-potent mouse hepatoblasts have been shown to preferentially differentiate into cholangiocytes at the periphery of circular micropatterned domains. Compared to central areas, these regions elicit increased actomyosin stress and elevate *NOTCH2* and *JAG1* transcriptional activity ^[42]^. Applying these principles to our system, we stipulate that rheological characterization of our optimized matrices will provide additional insight into the role of scaffold mechanics on Notch-mediated morphogenesis.

Aside from standardizing media components, differences in the described morphogenic potential of varying cholangiocytes should be reconciled by transcriptional profiling, which allows for stratification of specie specific phenotypes. In addition, single cell analysis can illuminate functional differences between stem cell-derived, adult progenitor and small/ large cholangiocyte subtypes ^[43]^. Finally, while immortalized cell lines have proven useful for *in vitro* mechanistic studies, questions regarding their phenotypic stability, tumorgenicity and transcriptional landscape remain, limiting their clinical utility for regenerative medicine applications. Our system provides an important framework for directing primary human biliary assembly that can be integrated with existing approaches in liver tissue engineering. There are notable limitations in this system, including the need for further functional validation of the cholangiocyte-like cells used in this study. For example, optimization of culture conditions should be conducted to limit mesenchymal features after serial passage and investigation of other markers such as the sodium-dependent bile acid transporter (ASBT) and osteopontin (OPN) should be pursued. With this described microfluidic system, one can measure cholangiocyte permeability, response to small molecules (ATP and acetylcholine stimulation on calcium influx dynamics), shear stress, and varying bile compositions. While we demonstrate the role of EGF stimulation on Notch signaling, other signal transduction pathways such as PI3K/Akt and MEK/ERK intracellular pathways can be affected by EGF as well. Furthermore, tethering EGF ligands to the scaffolds, rather than bulk administration, could lead to enhanced signal transduction and morphogenetic processes ^[44]^. In summary, we provide insight into growth factor-mediated branching in liver epithelium, consistent with findings in lung or kidney morphogenesis, but unveil tissue-specific ligand-receptor feedback between EGF and Notch signaling. Leveraging this insight, we directed the assembly of biliary duct structures using a microfluidic platform, adding a new model system of the portal triad.

## ACKNOWLEDGEMENTS

This work was supported by an NIH (EB008396) and the Howard Hughes Medical Institute (HHMI). Q.B.S is an HHMI Hanna Gray Fellow and S.N.B. is an HHMI investigator. We would like to specifically thank Dr. Heather Fleming for the critical editing of the manuscript. Dr. William Polacheck, Juliann Tefft, and Amy Stoddard provided technical support for the described microfluidic devices. We would also like to thank Dr. Jennifer Bays for performing the western blots, providing the CRISPR/Cas9 reagents, and giving technical assistance. Finally, we thank Dr. Amanda Chen, Dr. Tiffany Vo, Dr. Arnav Chhabra, and Keval Vyas for their technical advice and discussions throughout the project. This work is supported in part by the Koch Institute Support (core) Grant P30-CA14051 from the National Cancer Institute and the Koch Institute Swanson Biotechnology Center for technical support, specifically through the microscopy and flow cytometry core facilities.

## MATERIALS AND METHODS

### Primary Human Biliary Epithelial Cell Culture

Human intrahepatic biliary epithelial cells (IHCs) passage 1-5 (ScienCell, Carlsbad, CA) were cultured in complete epithelial growth media (ScienCell) on 50 µg/mL type I collagen coated surfaces (Corning). IHCs were isolated from human liver tissue using mechanical dissociation and enriched for CK-19. Media was exchanged every two to three days, passaged at 80% confluency with 0.05% trypsin/EDTA (ThermoFisher Scientific, Waltham, MA), and maintained in a humidified 5% CO_2_ incubator at 37°C.

### Flow Cytometry

Marker expression was verified through cytometry, where PE or APC-conjugated antibodies were stained on fixed and permeabilized cells. To harvest cells for flow analysis, serum was removed prior to adding TrypLE (Invitrogen, Waltham, MA) dissociation buffer, by washing with 1x PBS. After collection, cells were fixed with 3.7% paraformaldehyde (PFA) for 5minutes, spun for 2 minutes at 200xG, and resuspended in 0.1% Triton X for 10 minutes. Cells were then incubated in 100 µl of 0.1% bovine serum albumin (BSA; Sigma-Aldrich, St. Louis, MO) in PBS with conjugated antibodies for 1 hour at room temperature. Cells were washed three times to reduce non-specific staining and analyzed on a BD LSRFortessa (BD Biosciences, Franklin Lakes, NJ). To determine levels of expression, all analyses were conducted using IgG-PE or IgG-APC (BD) isotype controls.

### Immunofluorescence Staining and Imaging

Cells cultured on glass collagen I coverslips, were washed with 1X PBS and fixed in 3.7% paraformaldehyde for 10 minutes. After washing the samples with 1X PBS, samples were incubated in 0.1% Triton X-100 (Sigma-Aldrich) for 10 minutes. Next, cells were washed with 1X PBS and incubated for 1 hour at room temperature in 1% BSA to block for non-specific binding. Samples were then incubated with primary antibodies diluted in 1X overnight at 4°C, washed with incubated with secondary antibodies for 1 hour at room temperature. Finally, samples were incubated with Hoechst solution (ThermoFisher) for 3 minutes and washed with PBS prior to imaging. Cell morphology was identified using a Nikon TE200 Inverted microscope.

### Biochemical Assays

γ-Glutamyl Transferase activity was acquired using the Colorimetric Assay Kit (Sigma-Aldrich). In brief, 1 million, IHCs and fibroblast controls were collected in microcentrifuge tubes. Cells were pelted by spinning down at 200 x G for 3 minutes. After aspirating out the supernatant, cells were resuspended in 200 µl of cold GGT Assay Buffer and spun at 13,000 x G for 10 minutes. Activity was acquired through kinetic absorbance measurements (418 nm) at 37°C using a TECAN Infinite microplate reader. To determine alkaline phosphatase (ALP) activity, growth media of live cell cultures was removed, prior to washing vessels with pre-warmed DMEM/F-12. Adherent cells were then incubated for 20-30 minutes with a 1X ALP Live Stain solution (ThermoFisher). After incubation, the ALP solution was removed, and cultures were washed two times with fresh DMEM/F-12 for 5 minutes per wash prior to imaging using FITC illumination.

### Biliary Network Formation

Collagen gels were formed as previously described. In brief, 1.0×10^6 cells/mL were encapsulated in gels composed of Matrigel mixed at equal parts with 3.0 mg/mL rat tail collagen type I, that was titrated to pH 7.0-7.5 with 1M NaOH. 100 µl of the collagen mixture was added to wells of a 96 well plate and allowed to polymerize at 37°C for 30 minutes. After the gel solidified, an additional 100 µl of supplemented epithelial media was added.

### Biliary Network Quantification

A custom image processing program was written in MATLAB (Natick, MA) to quantify morphological features of branching phenotypes. Maximum intensity projections of confocal z-stacks were processed using ImageJ (NIH). Next, images were exported as JPEG files and imported into MATLAB. Red fluorescent protein (RFP) signal was used to identify network features. Masked image segments were enumerated and evaluated for percent coverage, branch length, and branch points.

### Quantitative Reverse-Transcription PCR

Total RNA was extracted using TRIzol Reagent (Invitrogen) from IHCs cultured under varying conditions. Quality and quantity of extracted RNA was verified by NanoDrop spectrophotometry prior to implementation of the 1 step RNA to Ct kit (ThermoFisher). Each measurement was conducted in triplicate with non-template controls using a BioRad CFX96 instrument. GAPDH served as endogenous controls for global normalization to acquire mRNA expression. Reference groups for differential analysis are outlined in the text and fold differences were calculated by the comparative Ct method.

### Lentiviral-Mediated CRISPR Genome Editing

CRISPR knockdown cells were generated using the lentiCRISPRv2 system (gift of F. Zheng, Addgene plasmid #52961). Scramble guideRNA (gRNA) (GCACTACCAGAGCTAACTCA) and NOTCH2 gRNA (GGCGCTCTGGCTGTGCTGCG) were designed using the Optimized CRISPR Design tool (F. Zheng, MIT) and cloned into the BsmBI site of plentiCRISPRv2. gRNA-containing pLentiCRISPR plasmids were co-transfected with pVSVG, pRSV-REV, and pMDL packaging plasmids into HEK-293T cells using calcium phosphate transfection. After 48 hours, viral supernatants were collected, concentrated using PEG-IT viral precipitator (SBI), and resuspended in PBS. Cells were transduced in growth medium overnight and selected with 2 μg/ml puromycin 48 h after infection. CRISPR modifications were verified by western blot.

### Western and Immunoblotting

Cell lysates were prepared with equal amounts of total protein (as measured using the Pierce Coomassie protein assay reagent) and separated on a NuPage Bis-Tris gels, transferred to PVDF (ThermoFisher), blocked in 5% milk and subjected to Western blot analysis using antibodies for Notch2 (Cell Signaling, D76A6) and GAPDH (Cell Signaling, D16H11). The blots were developed using ECL Western blot detection reagents (Pierce), and the signal was detected on iBrightTM CL1500 Imaging System (ThermoFisher).

### Quantification and Statistical Analysis

All statistical analysis was performed in GraphPad (Prism 9.0). Statistical significance was determined via methods outlined in the figure legends.

## SUPPLEMENTAL FIGURES

**Supplementary Figure 1.**
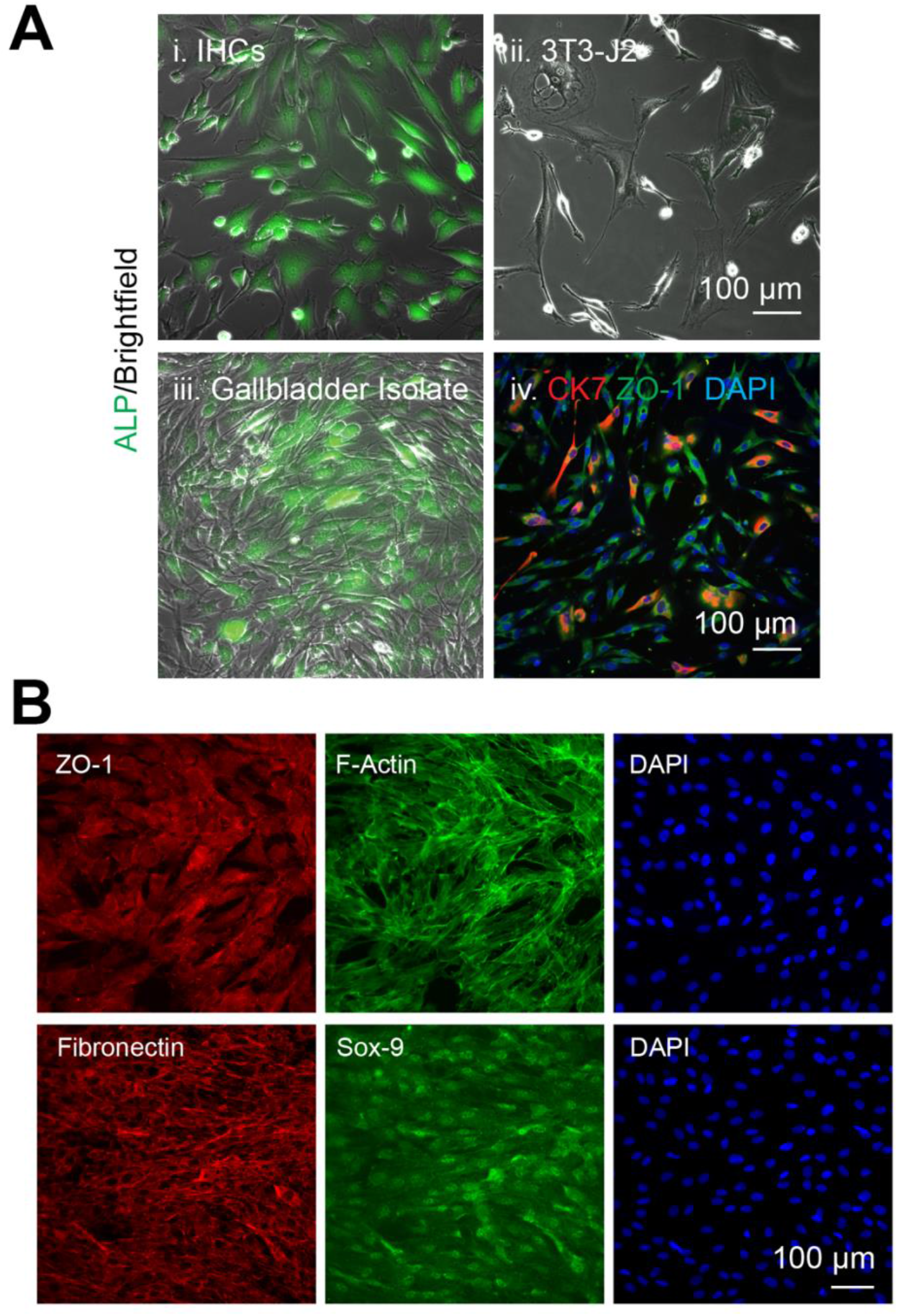
Characterization of Cholangiocyte Populations. (A) Alkaline phosphatase (ALP) uptake comparison between (i) intrahepatic cholangiocytes (IHCs), (ii) control J2-3T3 mouse fibroblasts and (iii) fresh extrahepatic cholangiocyte (EHCs) isolates from human gallbladder tissue. (iv) Immunofluorescence staining of fresh EHCs.

**Supplemental Figure 2.**
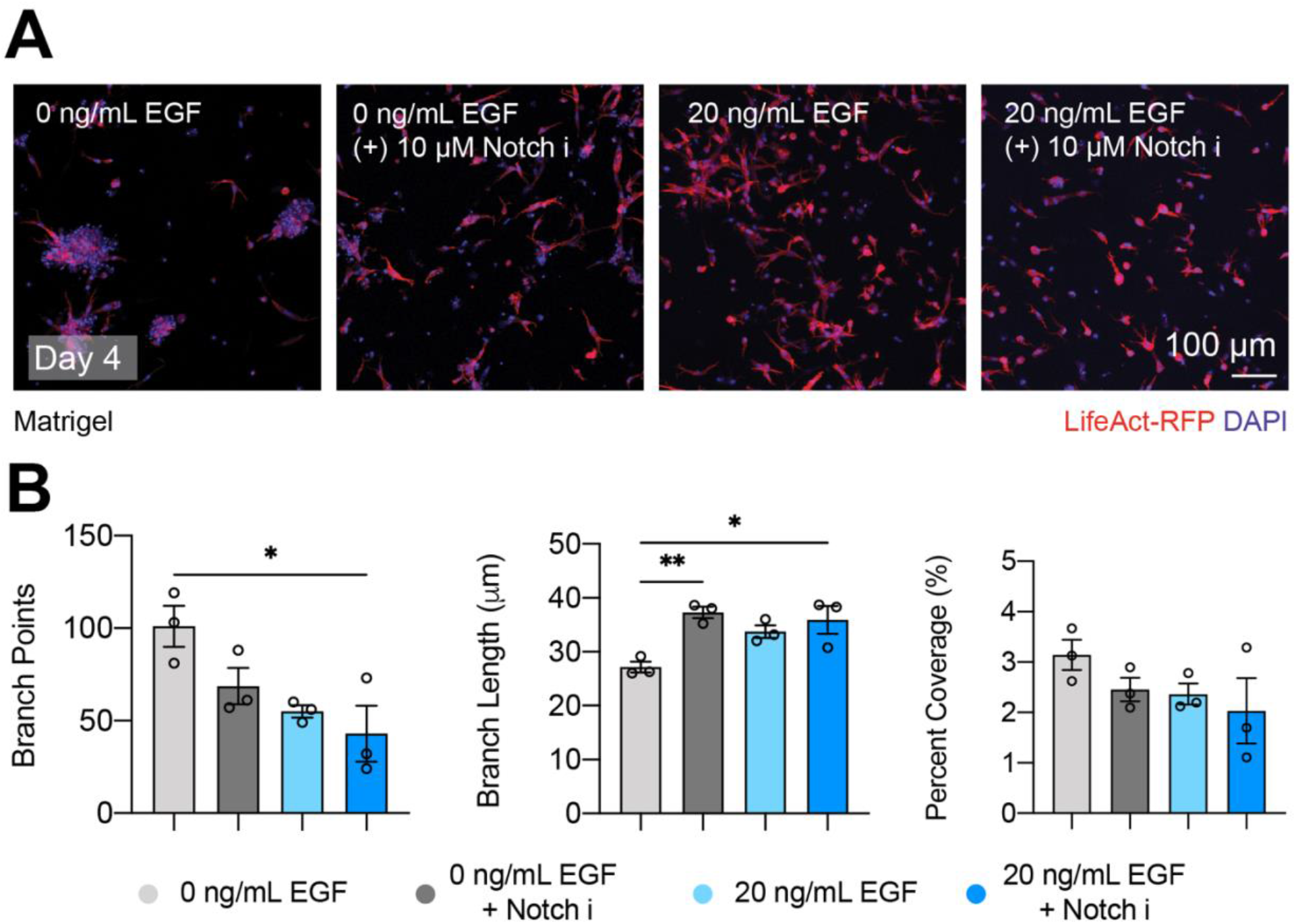
3D Culture in Matrigel Does Not Support Cholangiocyte Branching Morphogenesis. **(A)** Representative images of LifeAct RFP-IHCs encapsulated in Matrigel, cultured with and without EGF or Notch inhibition (10 µM L,685,458). **(B)** Quantification of branch length, points, and network percent coverage. Image data were generated from at least 3 independent fields of view from 3 biological replicate experiments. P-values were obtained by a One-Way ANOVA Tukey’s hypothesis test. P < 0.033 (*), P< 0.002 (**), P <0.001 (***). All data represented as mean ± SEM.

**Supplemental Figure 3.**
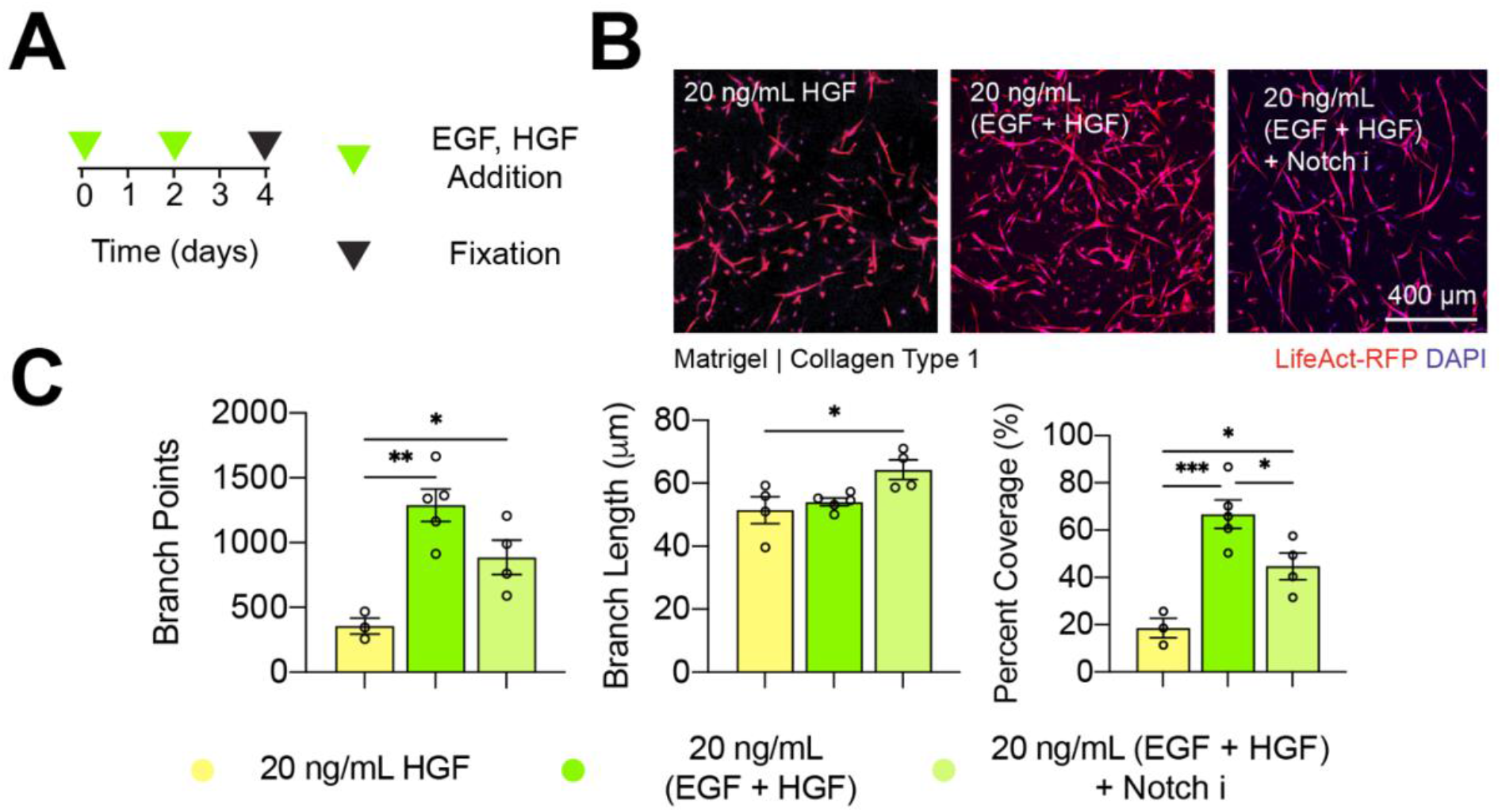
Hepatocyte Growth Factor (HGF) Supplementation Enhances Cholangiocyte Branching in Matrigel/Collagen Type 1 Hydrogel Blends. (A) Experimental approach to test the roles of HGF supplementation on IHC-RFP branching. (B) Representative maximum intensity projection of z-stack confocal images taken four days post encapsulation. (C) Quantification of percent coverage, major and minor axis length from at least 3 different fields of view from biological replicate experiments (n =3). P-values were obtained by a One-Way ANOVA Tukey’s hypothesis test. P < 0.033 (*), P< 0.002 (**), P <0.001 (***). All data represented as mean ± SEM.

**Supplemental Figure 4.**
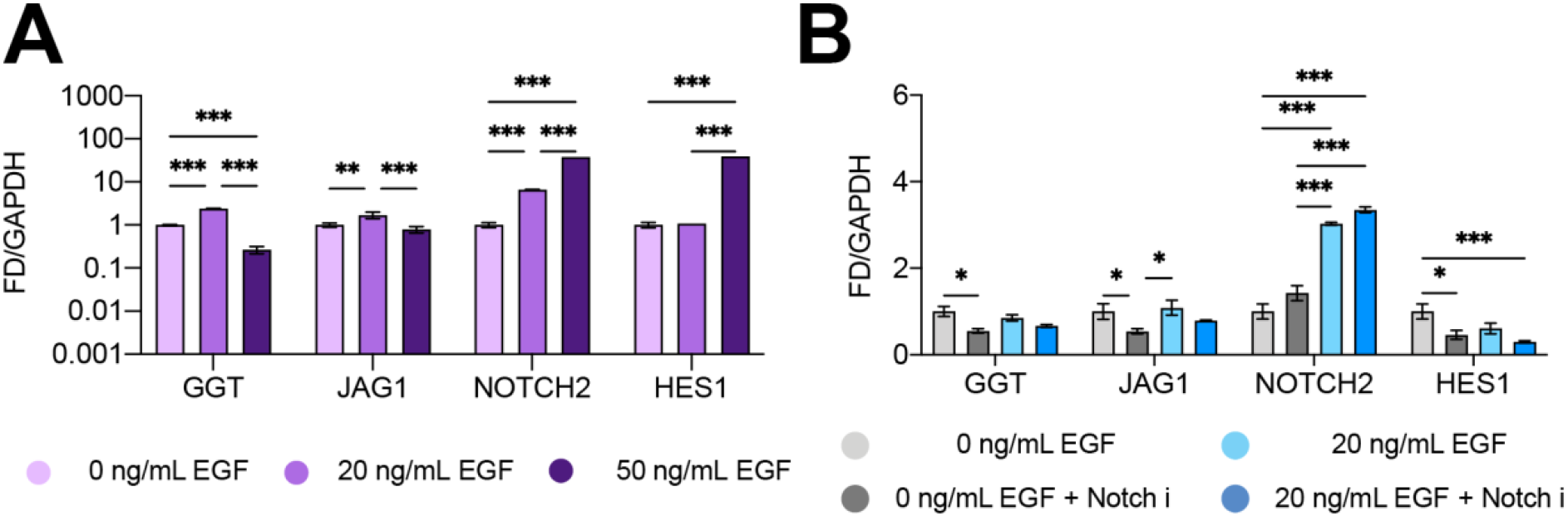
Effects of EGF Stimulation on Notch Signaling in 2D. (A) mRNA expression of *GGT, JAG1, NOTCH2* and *HES1* of IHCs cultured in the same dose regiment as 3D cultures (media exchanged every two days) with 0, 20 and 50 ng/mL EGF stimulation. Data is shown from pooled independent 2D cultures (n = 3 biological replicates). (B) mRNA expression of the aforementioned genes, of IHCs cultured in 2D, under 0 and 20 ng/mL EGF stimulation with and without Notch inhibition. Cells were harvested after 4 days following media exchanges every two days. P-values were obtained via Two-Way ANOVA Tukey’s hypothesis testing. P < 0.033 (*), P< 0.002 (**), P <0.001 (***). All data represented as mean ± SEM.

**Supplementary Figure 5:**
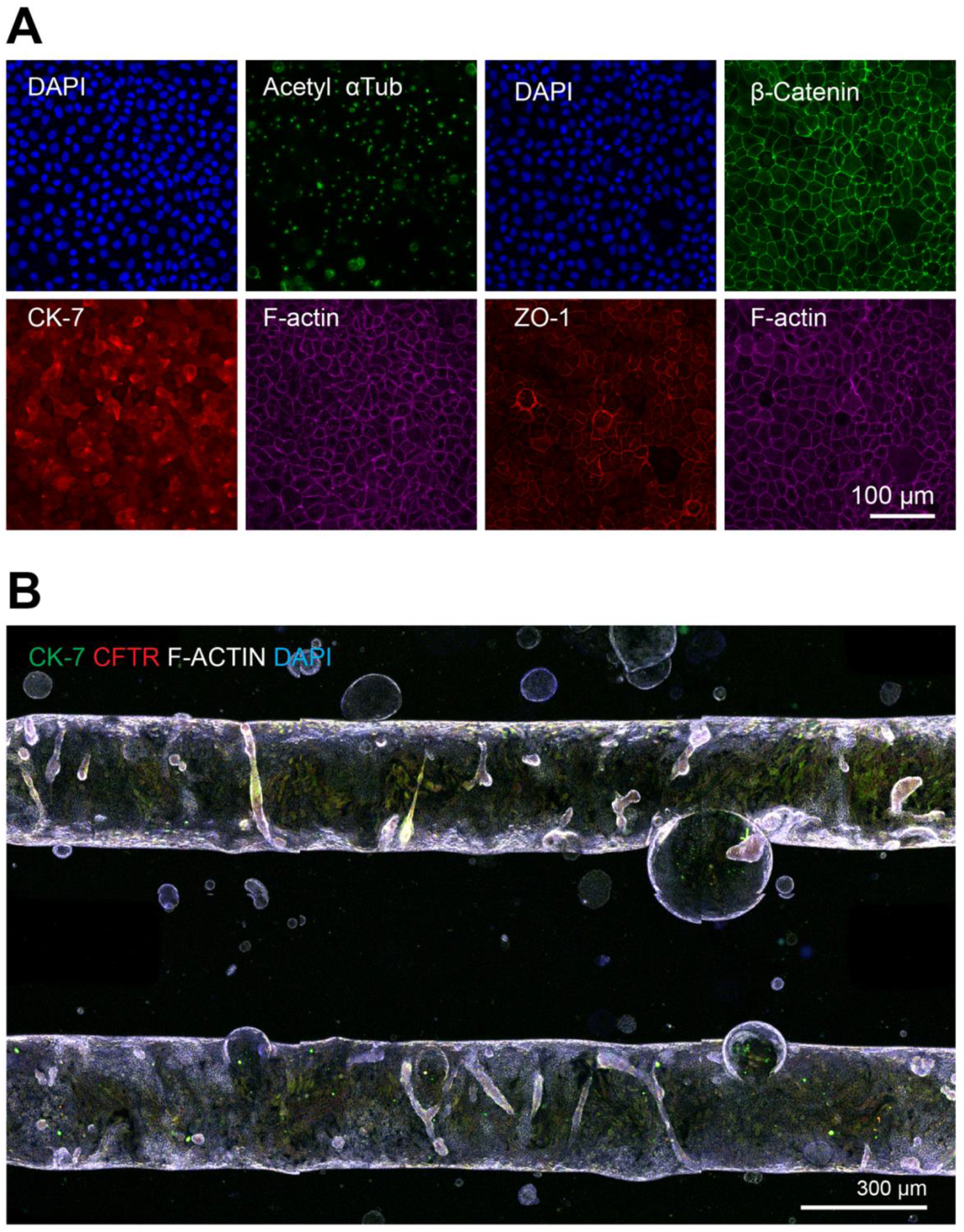
Microfluidic Culture of Normal Rat Cholangiocytes. (A) Immunofluorescence stains of normal rat cholangiocytes (NRCs) cultured in 2D for biliary (CK-7), primary cilia (acetylated α-tubulin; Acetyl αTub) and junctional markers (ZO-1 and β-catenin). (B) Microfluidic culture of NRCs in the dual microfluidic device.

**Supplementary Table 1: Primary and Secondary Antibodies**

**Supplementary Table 2: Reagents Used**

**Supplementary Table 3: q-RT PCR Primer List**

**Table S1:** Antibodies.

| Primary Antibodies | Source | Cat no | Host Species | Dilution |
| --- | --- | --- | --- | --- |
| Cytokeratin-19 | Abcam | ab7754 | Mouse | 1:200 |
| Cytokeratin-7 | Abcam | ab181598 | Rabbit | 1:100 |
| CFTR | Abcam | ab2784 | Mouse | 1:200 |
| Fibronectin | Abcam | ab2413 | Rabbit | 1:250 |
| ZO-1 | Invitrogen | 40-2200 | Rabbit | 1:100 |
| β-catenin | Cell Signaling | L54E2 (2677S) | Mouse | 1:5 |
| HNF-1β | Santa Cruz Biotechnology | sc-130407 | Mouse | 1:200 |
| SOX9 | Santa Cruz Biotechnology | sc-166505 | Mouse | 1:200 |
| Alexa Fluor 546 | Life Technologies (Eugene, OR) | A11010 | Goat anti-rabbit | 1:500 |
| Alexa Fluor 546 | Life Technologies | A11003 | Goat anti-mouse | 1:500 |
| Alexa Fluor 488 | Life Technologies | A21206 | Donkey anti-rabbit | 1:500 |
| Alexa Fluor 647 | Life Technologies | A22287 | Phalloidin | 1:1000 |
| Cytokeratin 7 Antibody (RCK105) PE | Santa Cruz Biotechnology | sc-23876 PE | Human | 1:200 |
| AFP Antibody (C3) Alexa Fluor® 647 | Santa Cruz Biotechnology | sc-8399 AF647 | Human | 1:200 |
| Anti-alpha 1 Antitrypsin antibody [EPR9090] (Alexa Fluor® 647) | Abcam | ab206735 | Human | 1:200 |
| PE Mouse Anti-Human IgG | BD Biosciences | 555787 | Human | 1:200 |
| Alexa Fluor® 647 Mouse IgG1 $\kappa$ Isotype Control | BD Biosciences | 557714 | | 1:200 |
| DAPI | ThermoFisher | D1306 |  | 1:50,000 |

**Table S3:**
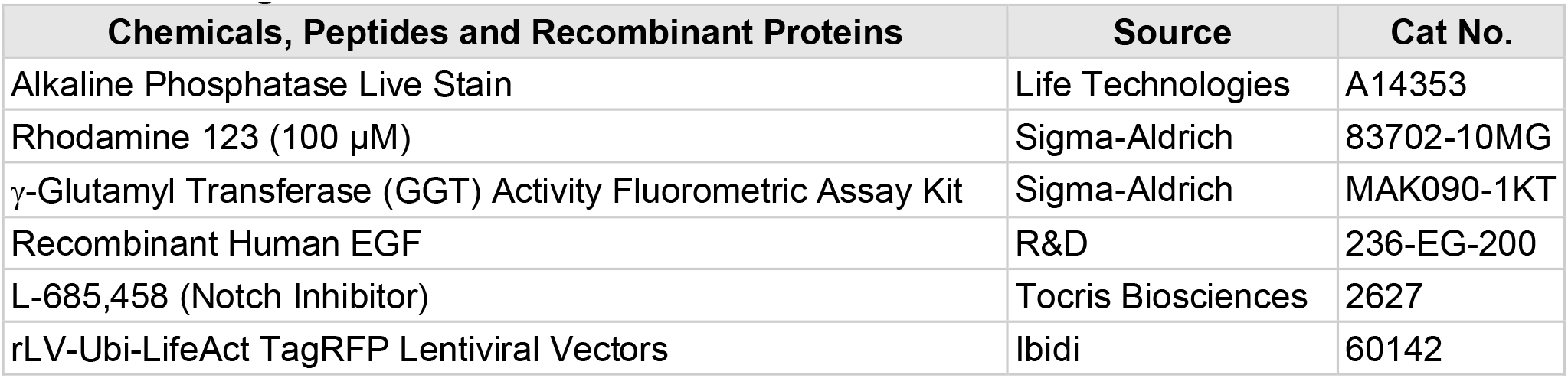
Reagents.

| Chemicals, Peptides and Recombinant Proteins | Source | Cat No. |
| --- | --- | --- |
| Alkaline Phosphatase Live Stain | Life Technologies | A14353 |
| Rhodamine 123 (100 µM) | Sigma-Aldrich | 83702-10MG |
| γ-Glutamyl Transferase (GGT) Activity Fluorometric Assay Kit | Sigma-Aldrich | MAK090-1KT |
| Recombinant Human EGF | R&D | 236-EG-200 |
| L-685,458 (Notch Inhibitor) | Tocris Biosciences | 2627 |
| rLV-Ubi-LifeAct TagRFP Lentiviral Vectors | Ibidi | 60142 |

**Table S2:** Primer Set.

| Primers | Source | Assay ID |
| --- | --- | --- |
| GGT | Life Technologies | Hs00980756_m1 FAM |
| JAGGED1 | Life Technologies | Hs01070032_m1 FAM |
| NOTCH2 | Life Technologies | Hs01050702_m1 FAM |
| HES1 | Life Technologies | Hs00172878_m1 FAM |

